## Supplementary material for "Personalized models of Disorders of Consciousness reveal complementary roles of connectivity and local parameters in diagnosis and prognosis": suplementary information

June 12, 2025

[1]Center for Brain and Cognition, University Pompeu Fabra, Barcelona, Spain
[2]Girona University, Spain
[3]Sorbonne Université, Institut du Cerveau - Paris Brain Institute - ICM, Inserm, CNRS, Paris, France
[4]Université Paris Cité, Paris, France
[5]Coma Science Group, GIGA-Consciousness, University of Liege, Belgium
[6]NeuroRehab & Consciousness Clinic, Neurology Department, University Hospital of Liège, Belgium
[7]Joint International Research Unit on Consciousness, CERVO Brain Research Centre, Laval University, Canada
[8]International Consciousness Science Institute, Hangzhou Normal University, Hangzhou, China

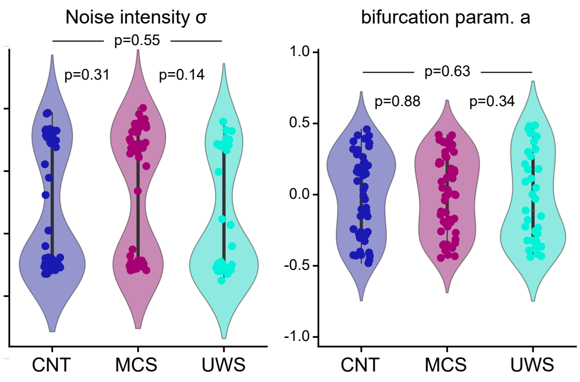


Figure S1: 1st level MBBs per condition obtained from the fitted node parameters: left, noise intensity $\sigma$, right, bifurcation parameter $a$, both dimensionless. Each dot is one patient.

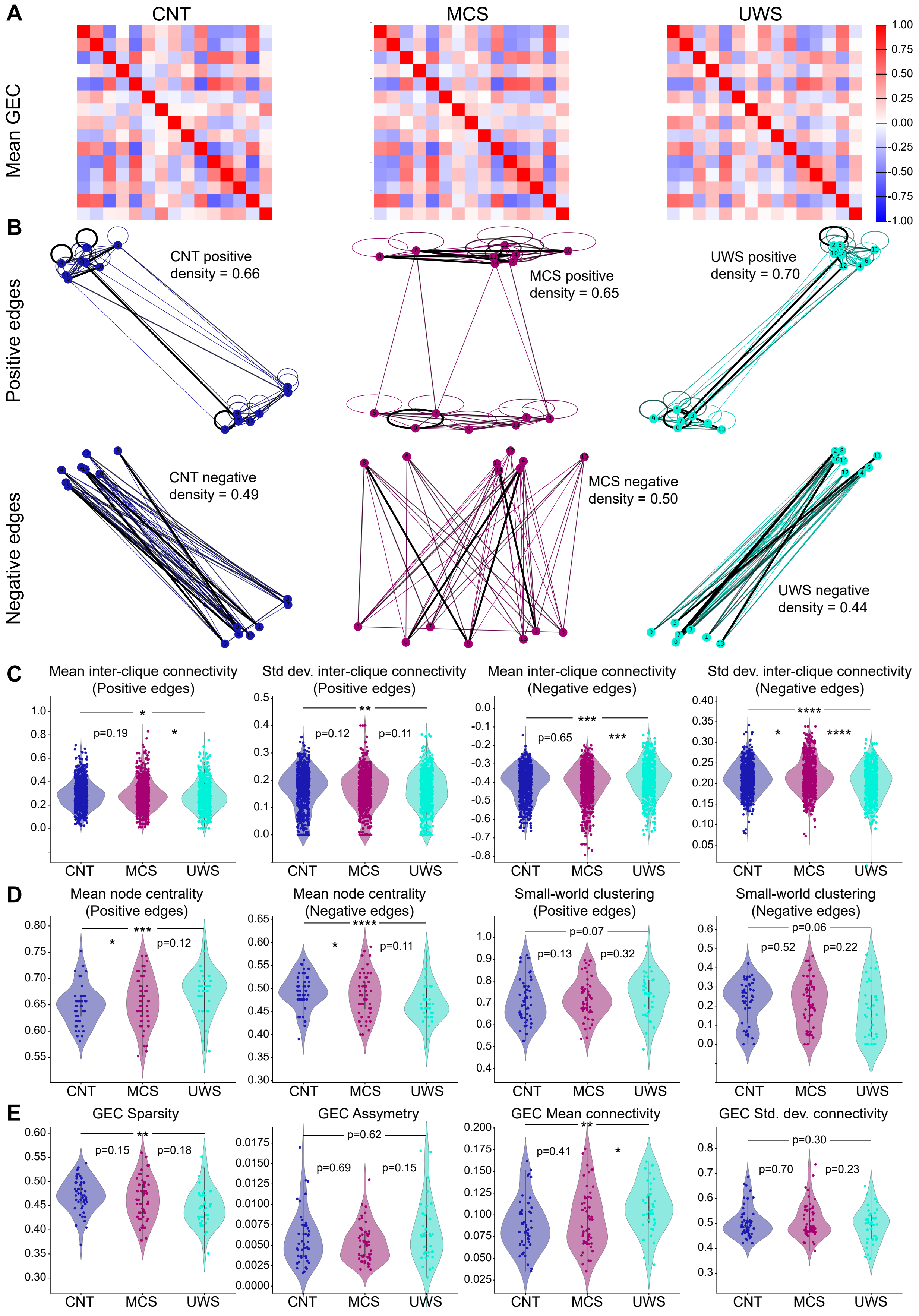


Figure S2: **GEC-based biomarkers for the Hopf model**: **A.** Average over all subjects of the fitted GEC matrix per condition (left to right CNT, MCS, UWS). **B.** Extracted graphs from the average GEC matrices (top: positive connections, bottom, negative connections) showing two subgroups (or cliques) of nodes positively connected between them and projecting negatively on the other subgroup, for each condition. **C.** Inter clique metrics: mean and standard deviation for positive (left) and negative (right) edges. **D.** Graph metrics: node centrality and Small-world clustering sigma parameter. E. GEC extracted measures: left to right: sparsity, asymmetry, mean and standard deviation values.

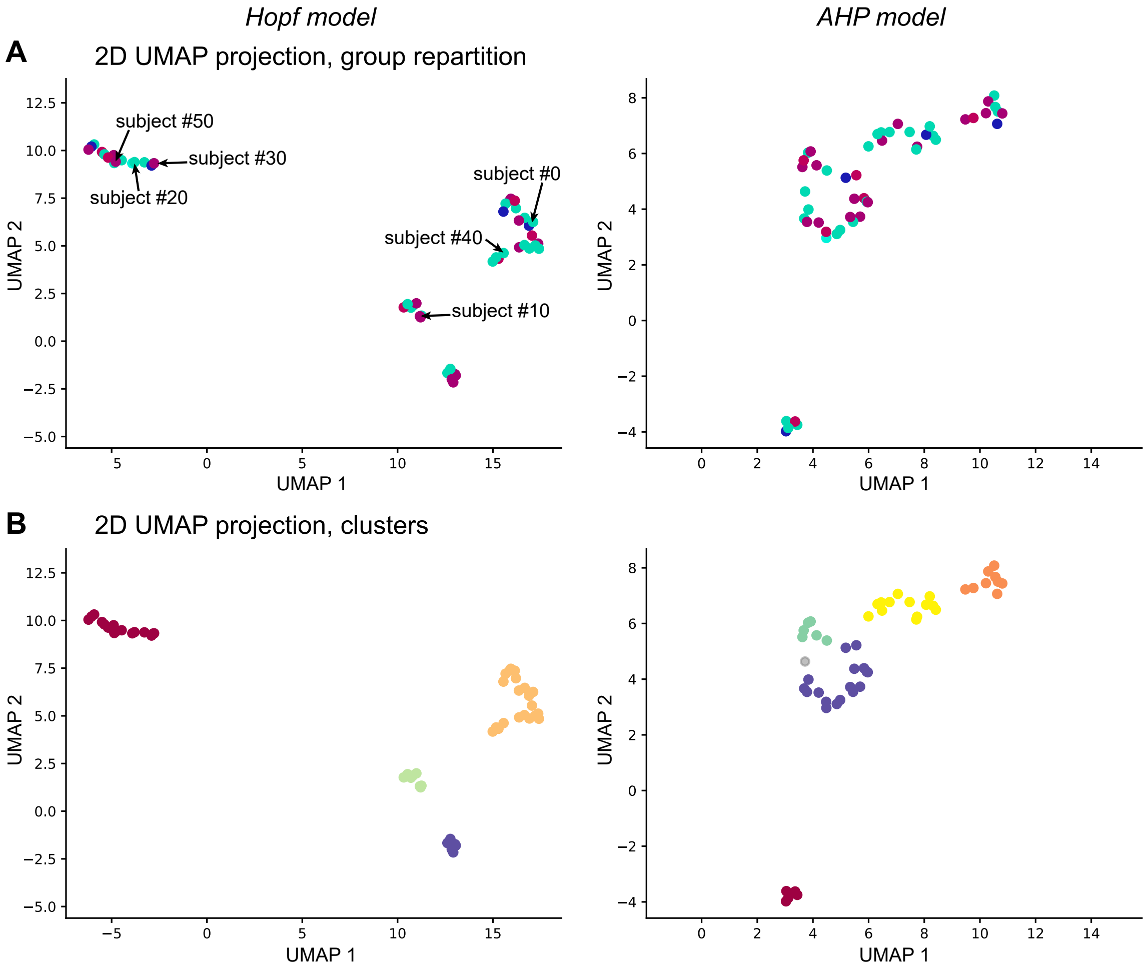


Figure S3: **UMAPs projections of the node MBBs**: **A.** 2D UMAP projection of the node MBBs (each dot corresponds to one patient), color coded by condition, for the Hopf (left) and AHP (right) models. **B.** same projection, color coded by cluster belonging, from the HDBSCAN clustering, for the Hopf (left) and AHP (right) models.

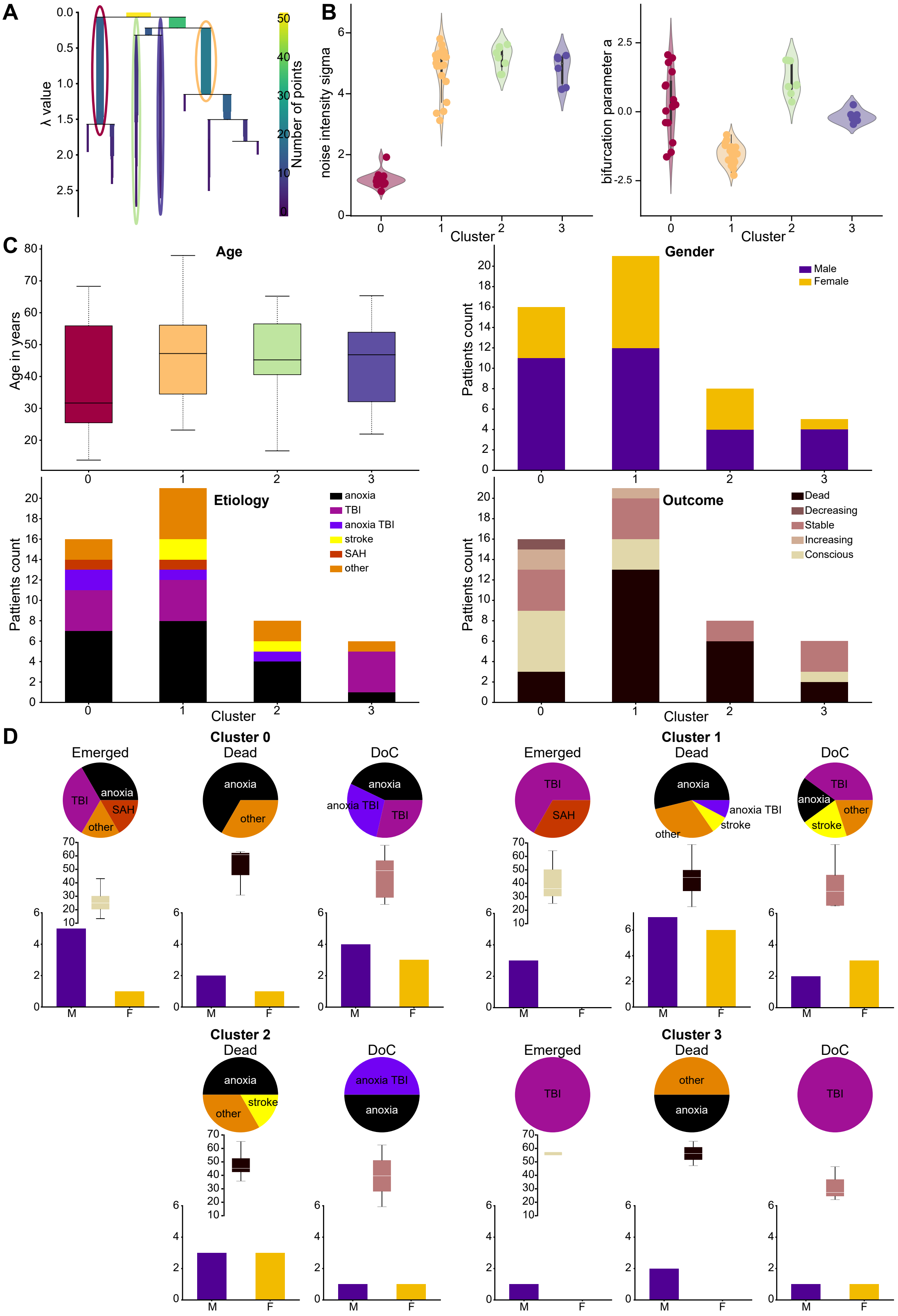


Figure S4: **Transverse clusters from the Hopf model node parameters**: bf A. Condensed tree plot of the HDBSCAN clustering algorithm showing the level of similarity between clusters. **B.** Fitted node parameters, separated by cluster, showing which parameters influence the attribution to which clusters. **C.** Composition of the clusters: top left, age distribution in years of the patients belonging to each cluster. Top right, gender distribution within clusters. Bottom left, etiology, bottom right, outcome. **D.** Co-factor analysis within clusters: for each cluster we split the patients between emerged, still in DoC, and dead and show the distribution of their etiology, age and gender balance.
